# Generalists versus specialists in fluctuating environments: a bet-hedging perspective

**DOI:** 10.1101/581371

**Authors:** Thomas Ray Haaland, Jonathan Wright, Irja Ida Ratikainen

**Affiliations:** Centre for Biodiversity Dynamics, Department of Biology, Norwegian University of Science and Technology (NTNU)

**Keywords:** Ecological specialization, environmental stochasticity, niche width, host specificity, performance curves, tolerance curves, canalization, plasticity

## Abstract

Bet-hedging evolves in fluctuating environments because long-term genotype success is determined by geometric mean fitness across generations. However, specialist versus generalist strategies are usually considered in terms of arithmetic mean fitness benefits to individuals, as in habitat or foraging preferences. We model how environmental variability affects phenotypic variation within and among individuals to maximize either long-term arithmetic versus geometric mean fitness. For traits with additive fitness effects within lifetimes (e.g. foraging-related traits), genotypes of similar generalists or diversified specialists perform equally well. However, if fitness effects are multiplicative within lifetimes (e.g. sequential survival probabilities), generalist individuals are always favored, since geometric mean fitness favors greater within-individual phenotypic variation than arithmetic mean fitness does. Interestingly, this conservative bet-hedging effect outcompetes diversifying bet-hedging. These results link behavioral and ecological specialization and earlier models of bet-hedging, and thus apply to a range of natural phenomena from habitat choice to host specificity in parasites.

**Impact summary:** Which factors determine whether it is better to be a specialist or a generalist? Environmental fluctuations are becoming larger and more unpredictable across the globe as a result of human-induced rapid environmental change. A key challenge of evolutionary biology is therefore to understand how organisms adapt to such variation within and among generations, and currently represents a knowledge gap in evolutionary theory. Here we focus on how traits evolve when the (changing) environment determines the optimal value of a trait, so that the optimal trait value changes unpredictably over time. Our mathematical model investigates how much variation is optimal in a trait. We expect specialists (low within-individual trait variation) to be favored in stable environments, with generalists (high trait variation) favored in more variable environments. We show that the answer depends on whether we look from the point of view of the individual or all individuals of the same genotype. If an individual does well in the short term, but its offspring all experience a different environment and therefore do badly, the genotype as a whole is in trouble, and will not be favored in the long term. One solution to this problem could be to produce offspring with different trait values, to ensure that at least some of the offspring do well no matter the environmental conditions they grow up in. This “don’t put all your eggs in one basket” diversification strategy is well-known in some organisms, but how helpful is it if there is also some within-individual (i.e. generalist) trait variation? By answering these questions under various environmental scenarios, we link together many different concepts in evolutionary ecology and animal behavior, increasing our understanding about how organisms may cope with the current changes in environmental conditions around the world.

## Introduction

Environmental fluctuations across different time scales pose a challenge for the evolution of organisms subjected to them, as well as for the researchers studying them (Botero *et al.* 2015; Tufto 2015). Notably, different fields of research have taken very different approaches to studying adaptations to stochastic environments, resulting in somewhat disparate bodies of literature dealing with quite similar topics (see Haaland *et al.* 2019).

Evolutionary and population biologists have long recognized that fitness effects across generations act multiplicatively (Dempster 1955; Cohen 1966; Frank 2011; Sæther & Engen 2015). In fluctuating environments, the optimal genotypic strategy is then the one that maximizes geometric mean fitness across environmental conditions (Lewontin & Cohen 1969; Simons 2002). Geometric mean fitness is vulnerable to variation in fitness. Thus, what appears suboptimal when considered from an individual’s point of view may in fact be optimal in the long term because it lowers genotype variance in fitness and thus increases geometric mean fitness across generations (Seger & Brockmann 1987; Simons 2002; Starrfelt & Kokko 2012). These bet-hedging traits (Slatkin 1974; Philippi & Seger 1989) have attracted considerable theoretical and empirical attention in recent years (Donaldson-Matasci *et al.* 2008; Olofsson *et al.* 2009; Childs *et al.* 2010; Simons 2011; Graham *et al.* 2014; Crowley *et al.* 2016), and are commonly split into two distinct types: diversifying and conservative bet-hedging (hereafter referred to as DBH and CBH, respectively). DBH strategies lower genotype variance in fitness by producing different offspring (Fig. 1A) and thus reduce the correlations in fitness among related individuals (Seger & Brockmann 1987; Hopper 1999; Einum & Fleming 2004; Starrfelt & Kokko 2012). In contrast, CBH strategies do not affect fitness correlation among individuals of the same genotype, but rather achieve lower variance in fitness for the genotype by lowering the variance in expected fitness for each individual separately. Such a CBH strategy often represents a generalist or ‘compromise’ phenotype that avoids doing badly in any one environment by doing moderately well in a range of environments, but gains lower arithmetic mean fitness across environments (Seger & Brockmann 1987; Starrfelt & Kokko 2012; Crowley *et al.* 2016).

**Figure 1:**
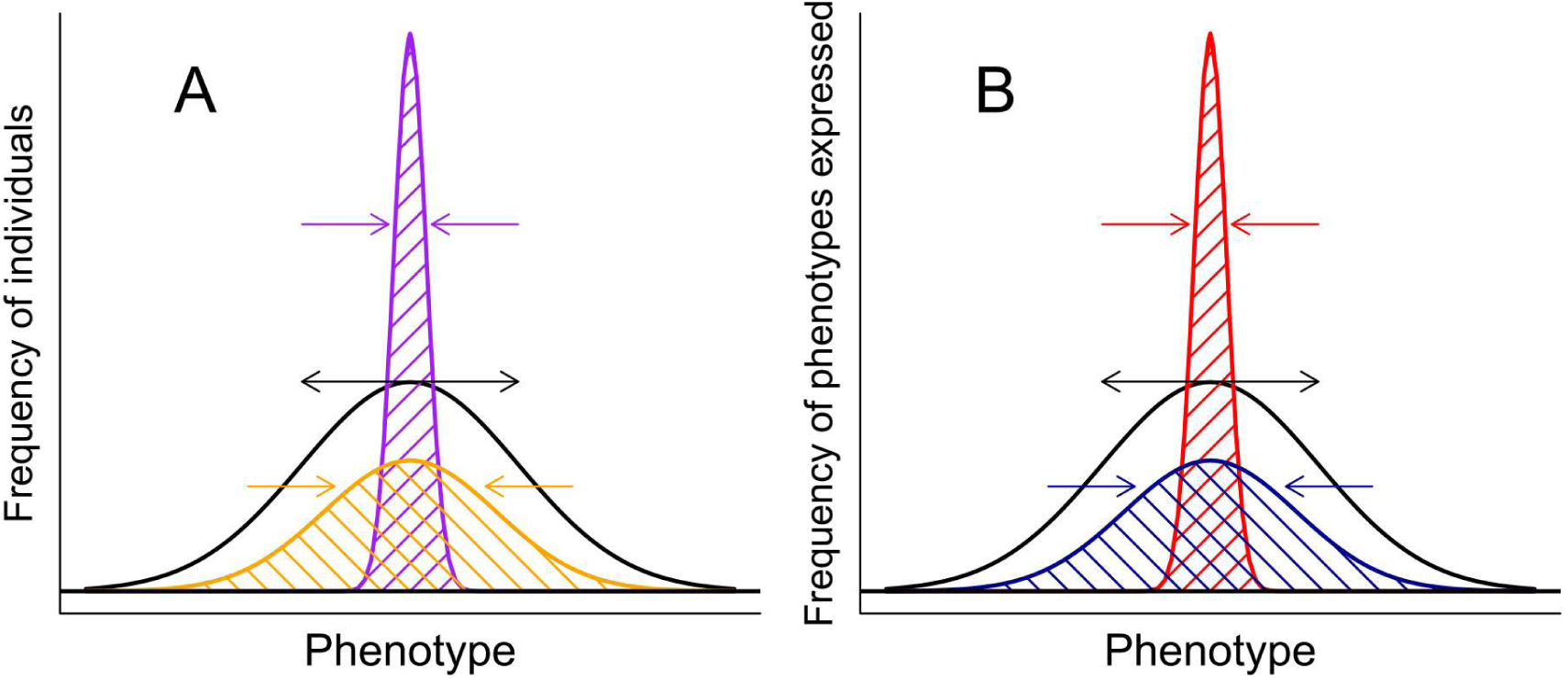
Sources of phenotypic variation among (A) and within (B) individuals. In both panels, the black curve represents the symmetrical fitness function (the position of which may fluctuate along the phenotype axis; fading black curves). In A, a genotype may produce a range of individual phenotypes with a narrow (purple curve) or wide distribution (orange curve). The colored arrows represent selective forces that can make the distribution of individuals narrower (purple; e.g. environmental canalization) or wider (orange; e.g. diversifying bet-hedging). In B, an individual may express a range of repeated phenotypes across events within its lifetime (e.g. behaviors or physiological performance) or across different parts of the organism (e.g. leaves or flowers) with a narrow (red curve) or wide frequency distribution (blue curve). In B the red arrows represent selection towards increased specialization (a taller, narrower distribution of phenotypes per individual), whereas the blue arrows represent selection towards increased generalization (a flatter, wider distribution of more different phenotypes per individual).

A related CBH strategy is ‘playing it safe’ in the face of an asymmetric fitness function, which provides an adaptive explanation for apparently suboptimal phenotypes seen in nature (Simons & Johnston 2003; Haaland *et al.* 2019). In their review, Starrfelt and Kokko (2012) claim that CBH and DBH are not mutually exclusive, but rather represent two ends of a continuum, and that both strategies can operate on the same trait. However, in a popular illustrative model considered in theirs and other bet-hedging papers (e.g. Hopper 1999; Crowley *et al.* 2016) it is not entirely clear how such a continuum should work. Haaland et al. (2019) have recently provided a solution to this issue via these ‘playing it safe’ CBH strategies in response to asymmetric fitness functions, which under certain conditions allow CBH and DBH to co-exist in the same trait as suggested by Starfelt & Kokko (2012). However, it is currently unclear how the original ‘compromise’ CBH strategy could coexist with DBH, as was originally suggested.

Largely parallel to this work on bet-hedging, other fields of research study adaptations to environmental variability from the perspective of the individual (i.e. maximizing some fitness proxy within a lifetime). In behavioral ecology, the optimal strategy or behavioral trait value is the one that maximizes expected (arithmetic) mean payoffs across all instances in which the trait is expressed over the course of a single lifetime (Maynard Smith 1982; Parker & Maynard Smith 1990; Westneat & Fox 2010), Despite this fundamental deviation from the bet-hedging view of evolution maximizing genotype geometric mean fitness, many topics in behavioral ecology echo the arguments of CBH and DBH while seeking to maximize individual arithmetic mean fitness. For example, insurance strategies in the face of an asymmetric fitness function due to starvation or predation risk (Brodin 2007; Dall 2010) have much in common with ‘compromise’ or ‘playing it safe’ CBH strategies (see Haaland *et al.* 2019), and generalist versus specialist foraging strategies as adaptations to within-lifetime fluctuations in prey availability (Dall & Cuthill 1997; Tinker *et al.* 2008) likewise focus on individual-level strategies and benefits (Fig. 1B) despite apparently mirroring the logic of DBH (Fig.1A). We note that such within-individual phenotypic variation can also be expressed across repeated parts of the individual in modular organisms, such as separate flowers or leaves of a plant, and the degree of this variation is known to be under strong genetic control (Diggle 2014).

Similarly, the field of thermal physiology also deals with adaptations to environmental variability, such as the width of thermal performance or tolerance curves (Lynch & Gabriel 1987; Gilchrist 1995) or apparently suboptimal body temperatures due to asymmetric fitness functions (Martin & Huey 2008; Angilletta 2009). While conceptually similar to more classical behavioral ecology studies, a key insight from these models is that the optimal strategy depends on whether fitness effects integrate additively or multiplicatively, within as well as across lifetimes. Additive fitness accumulation has been shown to favor specialist strategies even in highly variable environments, since high fitness gains in short periods where the organism is well suited to the environment can more than outweigh the long periods of no fitness gain when the environment fluctuates to unfavorable conditions (Gilchrist 1995). This scenario may be plausible for traits related to resource accumulation such as foraging strategies, locomotor ability or reproductive investment. In contrast, successive survival probabilities have multiplicative fitness effects. If environmental fluctuations away from the organism’s optimum are deadly, individual organisms simply seeking to maximize their own fitness (i.e. without any bet-hedging considerations) need to evolve a wider ‘generalist’ tolerance curve that spans the range of temperatures an individual can expect to experience within a lifetime (Lynch & Gabriel 1987; Buckley & Huey 2016) – see Fig. 1B.

Interestingly, fitness among individuals of the same genotype can also accumulate additively or multiplicatively. Typically this distinction is made between spatial and temporal environmental variability, or when referring to the ‘grain’ of the environment (Starrfelt & Kokko 2012). If environments only fluctuate spatially, such that individuals sharing a genotype experience different microenvironments within the same generation, but the mean environment is the same across generations (a fine-grained environment), genotype fitness is determined by the arithmetic mean fitness of all its bearers (McNamara 1995). Conversely, if environments only fluctuate temporally, such that all individuals in a generation experience the same environments and the environment fluctuates across generations (a coarse-grained environment), genotype fitness is determined by geometric mean fitness across generations. Thus, bet-hedging traits evolve in temporally varying environments to avoid low fitness in any one generation (Cohen 1966; Seger & Brockmann 1987; Venable & Brown 1988; Simons 2009; Starrfelt & Kokko 2012). Environments naturally vary both spatially and temporally, and the importance of bet-hedging scales directly with the relative magnitude of temporal versus spatial variability (Starrfelt & Kokko 2012; Gremer & Venable 2014; Crowley *et al.* 2016).

Thus, in order to link bet-hedging strategies with individual adaptations to environmental variation, we need to explicitly consider both within-and among-generation environmental variation, as well as both additive and multiplicative fitness effects within and among individuals. As Haaland et al. (2019) showed for insurance and ‘playing it safe’ CBH (i.e. with skewed fitness functions), traits maximizing arithmetic mean fitness in response to within-generation stochasticity may well reduce the scope for bet-hedging traits increasing geometric mean fitness in response to between-generation stochasticity. It is currently unknown how this interaction plays out between individual-level generalist-specialist strategies (Fig. 1B) and genotype-level bet-hedging strategies (Fig. 1A) with symmetrical fitness functions. Here we explore the potential for this type of generalist ‘compromise’ CBH strategy affecting the phenotypic ‘width’ of each individual during its lifetime, alongside the possibility of DBH affecting the width of the phenotypic distribution among individuals of the same genotype (Bolnick et al. 2007, see Fig. 1; Tinker et al. 2008). This comprehensive approach allows us to realize Starrfelt and Kokko’s (2012) suggestion of a continuum of bet-hedging strategies from CBH to DBH, while linking arguments for within-and among-individual phenotypic variation from a range of different research fields.

## Methods

We consider a continuous trait *z* with a symmetrical Gaussian fitness function *f*(*z*) that has a standard deviation of *σ_f_* and a mean or maximum fitness (optimum trait value) of *ϴ* that fluctuates due to environmental variation. We are then interested in the fate of a hypothetical genotype with three independent loci determining mean phenotype (*μ*), among-individual phenotypic variation (*σ*_a_), and within-individual phenotypic variation (*σ*_w_) - subscripts *a* and *w* here refer to among-versus within-individual variation, respectively. We begin by describing the fitness function 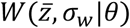 of an individual given its mean phenotype 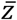 and phenotypic variation *σ*_w_ in a given environment *ϴ*.

### 1. Within-individual phenotypic variation

Within-individual phenotypic variation, *σ*_w_, is the trait determining the specialist/generalist continuum, and can be seen as the individual’s behavioral flexibility, variation across different parts of the individual (for modular organisms), or even degree of reversible phenotypic plasticity. While plasticity as such is not explicitly modelled here, we simply assume that a plastic individual is able to adjust its phenotype and thus gain some degree of fitness even if it is mismatched with the current environment (in that a higher *σ*_w_, creates a “flatter” fitness function – see Fig. 1B). Thus, our approach captures the outcome of this type of process in terms of the effect of within-lifetime variation in phenotypic expression on the subsequent fitness function. Regardless of the mechanism, we follow Lynch and Gabriel (1987) and model *σ*_w_ as the standard deviation of a normal distribution (which we call *g*_*w*_) around the individual’s mean phenotype 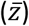. Such a definition ensures the traditional specialist/generalist trade-off, i.e. that the area under the phenotypic distribution is constant (and equal to 1), such that a wider distribution is necessarily lower at the mean than a narrower distribution, as in Fig. 1B (Kassen 2002; Phillips et al. 2014; but see Huey and Hertz 1984).

Thus, the fitness of an individual with phenotype *z* and phenotypic variation *σ*_w_ is the indefinite integral of the product of two normal distributions *f*(*z*)∼N(*ϴ, σ*_f_) and 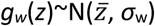. This integral can be computed and becomes a third Gaussian distribution with parameters that can be expressed as functions of the two input distributions (see Appendix A). Although both *f(z)* and *g*_*w*_(*z*) were probability density functions, i.e. they integrate to 1, 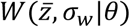 does not need to fulfill this criterion:

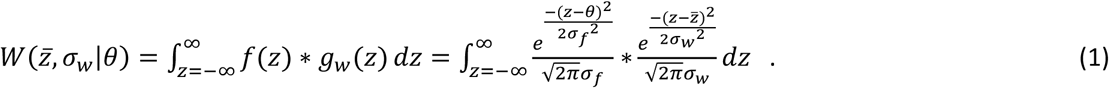

From this, we see how fitness depends upon the position *ϴ* of the fitness function relative to the position of the individual’s phenotypic distribution 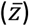. As we integrate over *z*, fitness contributions weigh more heavily when both factors provide larger values for the same *z*. Hence, having a phenotype that is well matched to the current environmental optimum provides largest fitness in all cases. However, the penalty for being mismatched (*ϴ*-*z*) becomes less severe for a larger *σ*_w_ i.e. for a generalist. Thus, generalists are less sensitive to fluctuations in the environment, and have a flattened fitness function compared to specialists, whose experienced fitness 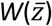 is close to *f*(*z*).

### 2. Among-individual phenotypic variation

We now add variability among individuals at the genotype-level, *σ_a_.* The gene value *σ_a_* can be seen as the degree of developmental instability, or the genotype’s sensitivity to stochastic microenvironmental fluctuations that determine an individual’s phenotype. Formally, we implement this variation similarly as for within-individual variation above, and let *σ*_a_ represent the genetically determined width of the Gaussian function around *μ.* The mean phenotype 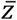 of each individual bearer of the genotype is therefore drawn from this distribution. A low *σ*_a_ represents environmental canalization of the phenotype, such that all bearers of the genotype will have phenotypes closely centered around *μ,* whereas a high *σ*_a_ would represent adaptive among-individual phenotypic variation (Fig. 1A), such as under diversifying bet-hedging (DBH).

Note that temporal and/or spatial environmental fluctuations among individuals within a generation can also drive the variation represented by *σ*_a_ (see Haaland et al. 2019). Therefore, a slightly different group of genotypic strategies, but with identical effect to the ones listed above, would be those modulating the degree of similarity versus difference between environments experienced by different individuals of the same genotype and phenotype. For example, niche-constructing traits such as building warmer nests to buffer against temperature fluctuations (Odling-Smee *et al.* 2003), active habitat selection based on genotype (Edelaar *et al.* 2008), or perhaps seeking more homogeneous habitats would all increase this similarity, whereas greater dispersal in space (or time, in the form of variation in dormancy) would decrease it. Thus, although our model ostensibly concerns the evolution of phenotypic variation within genotypes, it also has relevance for the evolution of various other traits that can affect the amount of environmental variation *experienced* within a genotype.

Whatever the mechanism, a genotype with mean phenotype *μ* and standard deviation among individuals in phenotype *σ*_a_ gains an average fitness 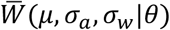, which is essentially the integral of the product of 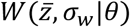 (eq. 1) and 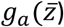, the distribution of individual mean phenotypes. This can be calculated using the formulae for the mean, variance and scaling constants of a Gaussian distribution resulting from the means and variances of the two other distributions (see Appendix A). We calculate 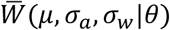 using numerical integration in R version 3.3.1 (R Core Team 2016) and the code is available on request.

### 3. Long-term genotype fitness in a fluctuating environment

The long-term fitness of a genotype in a fluctuating environment is typically approximated by the geometric mean fitness accumulated by the genotype across generations (Lewontin & Cohen 1969). Allowing the environmental optimum *ϴ* to fluctuate over time according to a distribution *h*(ϴ), long-term genotype fitness can be calculated by integrating 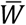 over the probability density of *ϴ*. Here we aim to specifically identify traits that fulfill the definition of bet-hedging - i.e. situations in which a given trait value decreases genotype arithmetic mean fitness and increases genotype geometric mean fitness. We therefore compare long-term arithmetic mean fitness across environments, which is given by

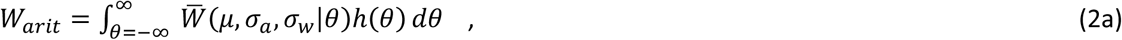

and long-term geometric mean fitness, which is the exponential of log 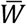 integrated across environmental conditions:

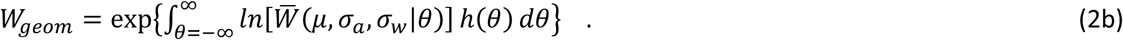

We calculate *W*_*arit*_ and *W*_*geom*_ for a range of combinations of *σ*_*a*_ and *σ_w_* in order to identify the optimal strategies of within-and among-individual phenotypic variation for different regimes of environmental fluctuations. We present results for scenarios where the environmental variation in *ϴ* follows a uniform distribution with different standard deviations *σ_ϴ_,* but also investigate scenarios where *ϴ* fluctuates according to a Gaussian distribution (see Supporting Information), to verify that the outcomes are not a consequence of the choice of *h*(ϴ).

### 4. Within-and between-generation temporal environmental fluctuations

To examine to what extent short-term individual-level strategies interact with or alter the scope for long-term genotype-level bet-hedging, we then explicitly add within-generation temporal environmental fluctuations, such that each individual experiences a range of environmental conditions over the period of its lifetime. When the environment fluctuates between generations (section 3 above), the genotype can maximize its long-term fitness either by adjusting the within-individual or the among-individual phenotypic width, or both. However, temporal fluctuations in the environment within a generation affect fitness accumulation within and among individuals quite differently. Importantly, if such within-individual environmental variation favors a more generalist phenotype at the individual level (larger *σ_w_*), then any bet-hedging effect in response to among-individual environmental variation would come about only if long-term geometric mean fitness favors a *different* (and presumably higher) level of ‘extra-generalist’ strategy as compared to that of long-term arithmetic mean fitness. We note that, though its effect is similar to that of diversifying bet-hedging (DBH) (Fig. 1A), such an extra-generalist strategy that increases *σ_w_* (within-individual phenotypic variation, Fig. 1B) beyond that which is adaptive in terms of maximizing arithmetic fitness would in fact constitute conservative bet-hedging (CBH), according to most of the literature (see Starrfelt & Kokko 2012). That is, it lowers genotype variance in fitness through lowering the variance in expected fitness for each individual, thereby maximizing geometric mean fitness at the cost of some arithmetic mean fitness. Thus, although the generalist may effectively ‘diversify’ its phenotype within its lifetime, any bet-hedging effect here of making individuals even more generalist would not constitute DBH, but rather CBH. However, diversification *among* individuals that lowers variance in fitness only at the genotype level would constitute DBH.

To explore these issues we assume that the microenvironment ϑ that a given individual experiences will fluctuate around the mean environmental condition *ϴ* for that generation according to the probability distribution *h(*ϑ|*ϴ*). Equivalent to eq. 1 for individual (lifetime) fitness, individual fitness in any one instance is now

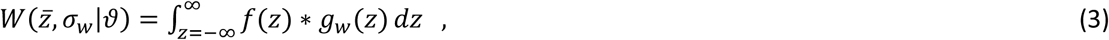

which depends upon the fitness function in the current microenvironment, *f*(*z*)∼N(ϑ, *σ_f_*), and the phenotypic distribution of the individual concerned, 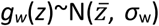. Then, to investigate the effects of additive versus multiplicative fitness accumulation within a lifetime (*sensu* ‘performance’ versus ‘tolerance’, c.f. Gilchrist 1995), we calculate either arithmetic or geometric mean fitness across the range of microenvironmental fluctuations, h(ϑ | *ϴ*). Using the same method as for eq. 2, we obtain for a given mean environment *ϴ*:

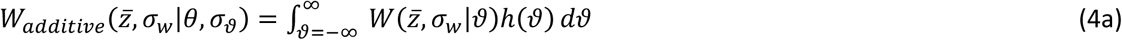

and

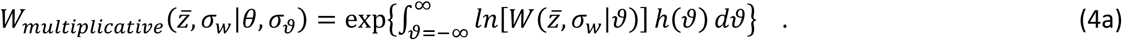

Eq. 4a and 4b can then be used instead of eq. 1 to calculate mean fitness when there is phenotypic variation within the genotype, following the steps in section 2 (above) and Appendix A. When the environment also fluctuates between generations, then long-term genotype arithmetic or geometric mean fitness can be calculated via numerical integration, as given in section 3 (above).

## Results

When there is no temporal environmental variation within generations (Figs. 2, S1), or when fitness effects within lifetimes are additive (Figs. S2, S7-8), extreme specialists are always favored, except when between-generation fluctuations *σ_ϴ_* are larger than the squared width of the fitness function (Bull 1987). This well-known result that is usually interpreted as the threshold above which among-individual phenotypic variation (i.e. DBH) becomes adaptive. However, contrary to previous treatments of DBH, the bottom row of Fig. 2 demonstrates that this adaptive phenotypic variation can arise both within and/or among individuals (i.e. via *σ*_w_ or *σ*_a_, respectively) with equal effect. This is indicated by the contour lines being always exactly symmetrical around the 1:1 diagonal in all panels (Fig. 2, bottom row). An increase in *σ*_w_, is adaptive in the rightmost panels because it lowers genotype variance in fitness across generations. This effect arises not because the correlations in fitness among individuals are lower (the variance-reducing mechanism created by DBH), but because each individual’s variance in fitness is lower, i.e. it is a CBH effect (see discussion above in Model Description, section 4). Thus, a genotype producing generalist individuals that have exactly similar phenotypes (CBH) does just as well as one producing specialist individuals that are highly diversified in their mean phenotypes (DBH). By comparing the top and bottom row (arithmetic versus geometric mean fitness landscapes) for the right-hand panels in Fig. 2 (*σ_ϴ_* >2), we clearly see that the greater geometric mean fitness gained by increased phenotypic variation comes at a cost of lower arithmetic mean fitness since arithmetic mean fitness is always highest at *σ*_a_ = 0 and *σ*_w_ = 0 (top row, Fig. 2), and thus these effects constitute bet-hedging.

**Figure 2:**
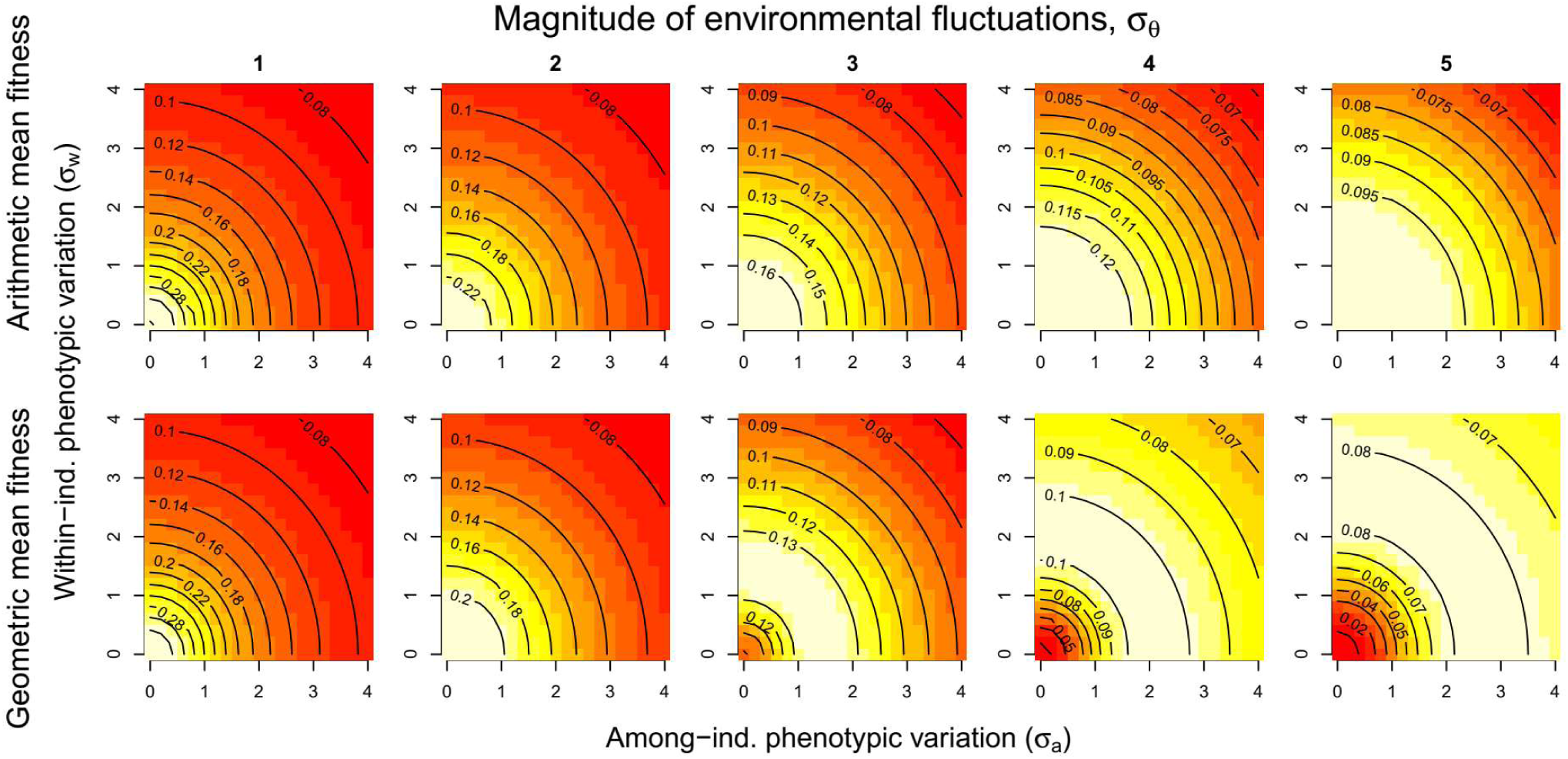
Fitness surfaces for genotypes with different amounts of among-(*σ*_*a*_; x-axis) and within-individual (*σ*_w_; y-axis) phenotypic variation. Contour lines show long-term arithmetic (top row) or geometric (bottom row) mean fitness in an environment where the phenotypic optimum *ϴ* varies across generations according to a uniform distribution around 0 with range ± *σ_ϴ_*, with values 1-5 across the different columns. (These *σ_ϴ_* values correspond to a standard deviation in *h*(*ϴ*) of 0.577, 1.155, 1.732, 2.309 and 2.887, respectively.) In this scenario there is no within-generation environmental variation (*σ*_ϑ_ = 0). Lighter background colors represent relatively higher fitness within each panel – note from the contour values that the different colors represent do not necessarily correspond to the same fitness values across panels.

We also note that, perhaps counter-intuitively, no matter how large the environmental fluctuations within or between generations, a genotype maximizing arithmetic mean fitness will always produce well-canalized specialists (i.e. all subplots in the top row of Fig. 2, S1, S7 and S8 have their fitness peak in the bottom left corner at *σ*_a_ = 0 and *σ*_w_ = 0).

However, when there are within-lifetime temporal fluctuations (*σ*_ϑ_ > 0) and fitness effects across instances are multiplicative, a different picture emerges. In Fig. 3, where *σ*_ϑ_ = 2 (see Figs. S2-S6 for a wider range of *σ*_ϑ_ values), both geometric and arithmetic mean fitness across generations favor the evolution of generalists (*σ*_w_ > 0). As *σ*_ϑ_ grows larger (Figs. S5 and S6), so does the amount of within-individual phenotypic variation *σ*_w_ that is favored. This arises from the fact that multiplicative fitness effects across instances within a lifetime will tend to favor individuals that are able to avoid low fitness values in any one instance, i.e. generalists. Such generalists are not following a bet-hedging strategy, but rather one that maximizes expected fitness of an individual (or arithmetic mean fitness across individuals).

**Figure 3:**
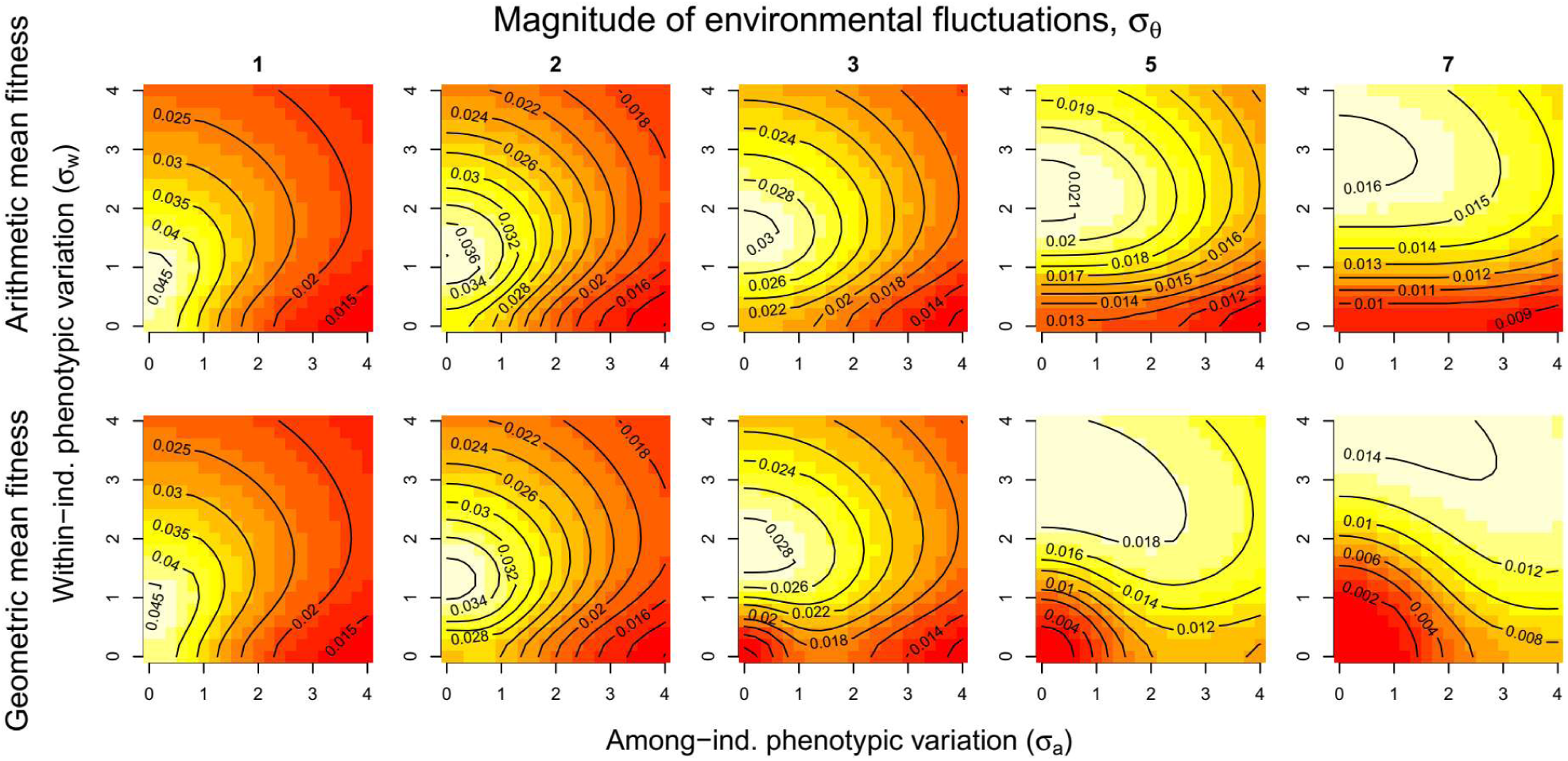
Fitness surface for genotypes with different amounts of among-(*σ_a_*; x-axis) and within-individual (*σ_w_*; y-axis) phenotypic variation when fitness effects are multiplicative. Contour lines show long-term arithmetic (top row) or geometric (bottom row) mean fitness in an environment where the mean optimal phenotype *ϴ* varies across generations according to a uniform distribution around 0 with range ± *σ_ϴ_* (with values stated above the different columns). Within generations experienced environments ϑ vary temporally (with multiplicative fitness effects) according to a uniform distribution around *ϴ* with range ± *σ*_ϑ_ = 2. Lighter background colors represent relatively higher fitness within each panel – note from the contour values that the different colors represent do not necessarily correspond to the same fitness values across panels.

As in Fig. 2, no adaptive among-individual phenotypic variation is expected in the top row of Fig.3, i.e. the peak of the fitness landscapes is always at *σ*_a_ = 0, and for any given value of *σ*_w_ the arithmetic mean fitness is always higher for lower *σ*_a_ (moving to the left on the fitness landscapes). We observe a similar pattern in the bottom row of Fig. 3 (and S2-S6), where within-individual phenotypic variation is favored but there is minimal adaptive among-individual phenotypic variation. Comparing the peaks of the arithmetic and geometric mean fitness landscapes (top versus bottom rows of Fig.3) reveals that geometric mean fitness favors considerable additional within-individual phenotypic variation on top of that already favored by arithmetic mean fitness (i.e. the peaks of the fitness landscapes in the bottom row are consistently higher up the y-axis than in the corresponding top row panel, see Table S1 for exact values). This ‘extra-generalist’ strategy again represents CBH – i.e. a genotype-level increase in geometric mean fitness at the cost of a lower arithmetic mean fitness, which comes about through lowering the fitness variance for each individual. Comparing the exact positions of fitness peaks when *σ*_ϑ_ = 0.5 versus *σ*_ϑ_ = 1 (Tables S2 and S3, respectively), we also note that increasing within-generation environmental fluctuations (*σ*_ϑ_) decreases the amount of this extra-generalist strategy attributable to CBH (rightmost columns), and this is because as *σ*_ϑ_ increases then the given amount of among-generation variation *σ_ϴ_* (i.e. the incentive for CBH) represents a smaller proportion of the total environmental variation.

In contrast to Fig.2, it matters in Fig.3 with its multiplicative fitness effects of events within lifetimes whether the phenotypic variation occurs within or among individuals. Specifically, no among-individual phenotypic variation *σ*_a_ is favored at the peaks of the geometric mean fitness landscapes in Fig. 3. However, the arithmetic and geometric mean fitness landscapes in Fig.3 differ notably in that there is a ridge of geometric mean fitness for higher *σ*_a_ when *σ*_w_ is lower than the peak (similar to that in Fig. 2), which does not appear for arithmetic mean fitness. Once the magnitude of between-generation environmental fluctuations in the optimum phenotype, *σ_ϴ_*, is larger than within-generation environmental fluctuations, *σ*_ϑ_ (i.e. at *σ*_ϑ_ = 2 in Fig. 3), a lineage with specialists (sub-optimally low *σ*_w_) will often benefit by increasing *σ*_a_. However, in all these cases increasing *σ*_w_ provides a steeper per-unit fitness gain than an increase in *σ*_a_. (To see this, start for example at *σ*_a_ = 0 and *σ*_w_ = 3 in the lower right panel in Fig. 3, and observe that the distance to the 0.014 fitness contour is much shorter in the *σ*_w_ (y-axis) direction than in the *σ*_a_ (x-axis) direction.) See also Table S2 for a possible small exception.

## Discussion

Broadly, our results exploring additive (Figure 2, S1) versus multiplicative (Figure 3, S2-S6) fitness effects within lifetimes support the results found in previous theoretical work. For example, Lynch and Gabriel (1987) conclude that with multiplicative fitness effects within an organism’s lifetime the optimal tolerance curve (see Introduction) must be wide enough to span the range of temperature fluctuations experienced within a lifetime, whereas Gilchrist (1995) uses additive fitness effects within lifetimes and finds that narrow performance curves are optimal in most environments. A recent simulation study combining both additive and multiplicative fitness components to investigate the evolution of thermal niches in *Drosophila* was able to successfully predict a latitudinal gradient in thermal tolerance (Buckley & Huey 2016). Our model should provide a more general framework within which to place future work of this type on specific systems showing such adaptations.

Our modeling framework has numerous applications beyond thermal physiology. In particular, if within-individual phenotypic variance represents distributions of behaviors (specialists always choosing the same option, versus generalists choosing a range of different options) we can interpret our results in a range of contexts of interest in behavioral ecology and evolutionary ecology, such as foraging behavior (e.g. optimal diet breadth, Dall & Cuthill 1997), reproductive decisions (e.g. timing of reproduction, Nussey *et al.* 2005; nest site placement, Bailey *et al.* 2017; size vs number of offspring, Ratikainen *et al.* 2018), or host-parasite interactions (inducible herbivore defenses, Van Tienderen 1991; host preference in brood parasites, Feeney *et al.* 2014). These topics differ in which of the model variants presented here would be most relevant. For example, for traits related to reproductive decisions, an important distinction would be between semelparous and iteroparous organisms. In organisms that just reproduce once per lifetime, traits affecting reproductive decisions only experience multiplicative fitness effects across generations. However, in organisms breeding several times throughout their lifetime, lifetime reproductive success can be determined by the sum of the reproductive output in each breeding attempt (additive effects).

Conservative bet-hedging (CBH) has often been conceived in the form of generalist individuals able to perform moderately well across a range of environmental conditions (a ‘compromise’ phenotype, see Introduction). However, such CBH does not typically outperform diversifying bet-hedging (DBH), if DBH produces specialists that perform well in their preferred environment and in the proportions that various environments are likely to occur. In the classic bet-hedging model of reproduction in dry versus wet years (Seger & Brockmann 1987; Starrfelt & Kokko 2012), this clearly changes if we include survival across many years before an individual is able to reproduce, the optimal strategy in the long run must be to be able to tolerate both dry and wet conditions. Bet-hedging is obviously not needed to explain generalist adaptations to such within-lifetime or ‘fine-grained’ temporal variation among discrete environmental conditions (Seger & Brockmann 1987; Reboud & Bell 1997; Kassen 2002). However, by generalizing this to a model with continuous environments we are able to not only show the selective advantage of generalists over diversified specialists, but also that a CBH component favors even wider within-individual phenotypic distributions (‘extra-generalists’) than the individual’s optimum phenotypic width, and that this appears more effective than any DBH strategy in nearly all cases (Fig. 3). Thus, our model suggests that the scope of adaptive DBH in nature may be limited to situations where generalist phenotypes are somehow constrained and/or are not able to ensure survival under the full range of environmental conditions experienced within a lifetime, or where fitness effects are only additive within lifetimes. In these latter cases, genotypes consisting of canalized generalists and diversified specialists perform equally well.

Foraging theory is a classical field of behavioral ecology where the use of arithmetic average pay-offs (e.g. net energy gains) have long been used to successfully predict optimal diet widths (Futuyma & Moreno 1988; Stephens *et al.* 2007), and especially to explain the observed tendency towards diet specialization across a range of taxa (Forister *et al.* 2015). Recent work has, however, revealed that even in generalist species there exists considerable variation among individuals in diet choice (and other traits), with each individual specializing on a much more restricted range of prey types than the population as a whole (Estes *et al.* 2003; Bolnick *et al.* 2007; Dall *et al.* 2012). The question of whether more generalist populations should also contain more among-individual variation (the ‘niche variation hypothesis’) has received both empirical and theoretical support, but also criticism (see Bolnick *et al.* 2007). In the context of our model, for foraging-related traits with strictly additive fitness benefits across foraging bouts, we predict specialization up until the threshold amount of environmental fluctuations given by Bull (1987), after which symmetrical combinations of among-and within-individual phenotypic variation are expected to be equally viable in the long term (Fig. 2, bottom right panels). In other words, our very general model predicts that if the distribution of available prey types is sufficiently variable in time, a population consisting of different specialists (spanning the range of prey types) is as likely to evolve as a population of less differentiated generalists (with individual diet width spanning the same range of prey types), all else being equal. Indeed, a study of Brünnich’s guillemots (*Uria lomvia*) found large individual differences in both diet composition and degree of diet specialization, but across 15 years there was no difference in fitness components (survival or reproductive success) between specialists and generalists (Woo *et al.* 2008). This is consistent with the analytical model predictions we present here, which have also been confirmed using simulation models (Haaland, unpublished results). Along this ridge of equal fitness (the diagonal in Fig.2), we also predict that an increase in within-or among-individual diet width (cf. Tinker et *al.* 2008) necessarily requires a decrease in the other component. However, we acknowledge that for simplicity and generality our model obviously omits several factors commonly involved in explanations of individual specialization and observations of character displacement, most notably intraspecific competition and density-and frequency-dependent effects resulting from resource depletion (Svanbäck & Persson 2004; Tinker *et al.* 2012), which may have important additional effects.

Our predicted continuum of among-versus within-individual variation in resource use also interestingly shows up in the multiple solutions in host preference of different species of brood parasites (Davies 2000). For example, most cuckoos are host generalists at the species level, but populations consist of segregated ‘gentes’, each specialized on a different single host species and with special adaptations such as egg size and coloration closely matching that of its preferred host (Brooke & Davies 1988). Availability of suitable hosts varies spatially and temporally, due not only to fluctuations in host population sizes, but in some species also because the host species have evolved highly variable egg phenotypes as a defense against brood parasitism (Spottiswoode & Stevens 2011). Hence, highly specialized gentes are always vulnerable to a shortage of suitable hosts, and any genotype that hedges it bets will do better in the long run. Indeed, brood parasites specialized on a single host will consistently lay some eggs in the nests of ‘wrong’ hosts (Davies 2000, chapter 9). Although these eggs will often fail, occasionally they may be successful and lead to daughters imprinted on the new host species (Moksnes & Røskaft 1995; Davies 2000), thus creating heritable (non-genetic) phenotypic variation. This might therefore be seen as a diversifying bet-hedging (DBH) strategy favoring the long-term survival of a genotype despite sacrificing some reproductive success in the short term. Theoretically, whether such DBH becomes adaptive depends upon the temporal fluctuations in the environment (*ϴ* above) among relative to within generations (Figs. S7-S8), and relative to the width of the fitness function, *σ_f_* above (Bull 1987; Botero *et al.* 2015; Haaland *et al.* 2019). Thus, a narrow fitness function (e.g. high egg specialization, leading to poor viability if parasitizing the wrong host) and fluctuations in abundance of host species can adaptively favor these consistent ‘mistakes’ and strong offspring imprinting in many European cuckoos. In contrast, cowbirds or species such as the generalist brood parasite Horsfield’s bronze-cuckoo (*Chrysococcyx basalis*) have eggs that do not mimic a single host exactly, and there is no need for offspring to preferentially lay eggs in the nests of the same species in which they were raised (Feeney et *al.* 2014). Furthermore, since Horsfield’s bronze cuckoo is a nomadic, opportunistic species breeding year-round wherever conditions are favorable, the within-generation fluctuations in host availability likely far outweigh the among-generation fluctuations, also favoring a lower *σ*_a_.

It has been suggested that environmental variation can affect the host specificity of parasites in other systems as well, including avian malaria parasites *Plasmodium* and *Haemoproteus* spp. (Fecchio *et al.* 2019). The short life cycles of vector-borne pathogens will lead them to experience environmental variation (host availability) mostly at the among-generation level. While an in-depth study of the life cycles of various pathogens is beyond the scope of this paper, we suggest that useful starting points in identifying potential generalist parasites may be identifying bottlenecks in the availability of hosts at different life stages, e.g. through time series data of population trends. If the availability of specific hosts fluctuates more (i.e. large *σ_ϴ_*), it is likely that evolution has favored either more diversified parasite strains, or less host-specific parasites.

In summary, we have shown how environmental fluctuations within and among generations affect the optimal amounts of phenotypic variation within and among individuals to maximize long-term genotype fitness. Our model identifies additive versus multiplicative fitness effects within lifetimes as a critical factor in determining the evolution of specialist versus generalist phenotypes. It shows that when generalists are favored, CBH via increasing within-individual phenotypic variation (‘extra-generalist’) outperforms DBH that increases only among-individual phenotypic variation. Our results have general implications for understanding the relative importance of generalist-specialist strategies and the potential coexistence of the different bet-hedging strategies (CBH versus DBH), and provide novel links with well-developed fields such as thermal physiology, behavioral ecology and host-parasite interactions. While additive versus multiplicative fitness effects have been discussed in light of thermal or environmental ‘tolerance’ (Lynch & Gabriel 1987; Gilchrist 1995; Buckley & Huey 2016), how these contrast and interact has not previously been fully integrated with bet-hedging theory and the surrounding literature, or with generalist-specialist research within behavioral ecology. We hope that our model serves to clarify these findings amidst a growing interest in bet-hedging, phenotypic plasticity and evolutionary adaptations to anthropogenic environmental change. Finally, we note that the relative importance of additive versus multiplicative fitness accumulation among and within individuals is itself a trait that can be subject to selection, for example through the evolution of lifespan (see Ratikainen & Kokko 2019), semelparity versus iteroparity and pace-of-life syndromes (Wright *et al.* 2019), and that altered selection pressures from these changes can feed back on types of phenotypic evolution we explore here. Therefore, what appear in our model as assumptions and parameter values may well be traits that have coevolved alongside the generalist-specialist and bet-hedging strategies that we explore, suggesting the need for further theoretical investigation of this issue.

## Authorship

All authors developed the ideas. TRH built the model. All authors analyzed the results. TRH wrote the manuscript with input from all authors.

## Data accessibility

R code is available on request.

## Acknowledgements

We thank Jarle Tufto for help with the R code and Gunnar A. Sveinsson for mathematical advice. TRH and IIR are supported by the Research Council of Norway on grant 240008 awarded to IIR on the Young Talented Researchers program, and this work was partly through its Centres of Excellence funding scheme, project number 223257, to Centre for Biodiversity Dynamics (CBD) at the Norwegian University of Science and Technology (NTNU). The authors declare no conflicts of interest.

## Appendix A: Calculating fitness functions under phenotypic variability

The product of two Gaussian distributions with means *a* and *b* and standard deviations *A* and *B,* is a Gaussian distribution with mean 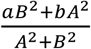, standard deviation 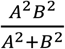, and multiplied with the constant 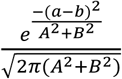. Using these formulae, we can calculate the fitness of an individual with phenotypic distribution 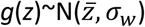 in an environment where the fitness function is *f*(*z*)∼N(*ϴ, σ_f_*), which is the indefinite integral of the product of these two Gaussian distributions. Setting *a=ϴ, A=σ_f_* (mean and sd of the first multiplicand, the fitness function 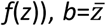 and *B=σ_w_* (mean and sd of the second multiplicand, the phenotypic distribution *g*(*z*)), we find that

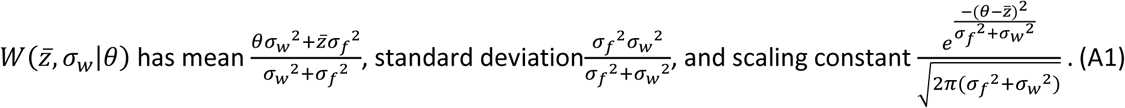

These values can be plugged into equation 1 (main text), which then gives the complete form of the fitness function 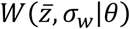:

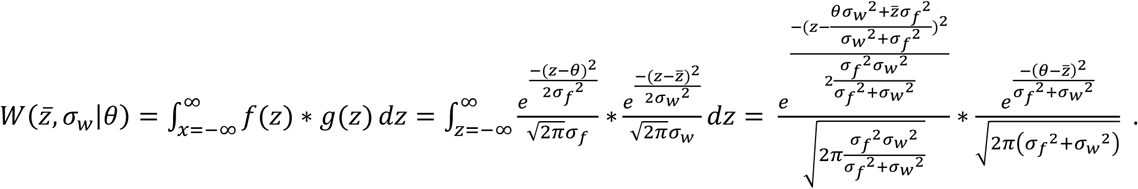

We use the same method to calculate average fitness of a genotype with mean phenotype *μ,* among-individual variance *σ_a_^2^*, and within-individual variance *a*_*w*_^2^, 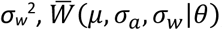, which is found by integrating the product of 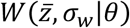 and *g*_*a*_(*z*). We do not show the resulting 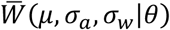 here, since it becomes very involved, but using the same formulae as above it can be calculated by plugging in 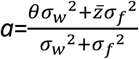, i.e. the mean of 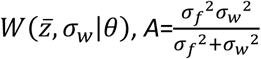, the standard deviation of 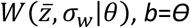, *b=ϴ*, and *A*=*σ_ϴ_*.

