## Supplementary Information for "Generalists versus specialists in fluctuating environments: a bet-hedging perspective"

To

### Contents

2. Figure S1: Fitness surfaces with Gaussian $h(\Theta)$

3. Figure S2: Fitness surfaces with $\sigma_\vartheta$ = 0.5

4. Figure S3: Fitness surfaces with $\sigma_\vartheta$ = 1

5. Figure S4: Fitness surfaces with $\sigma_\vartheta$ = 1.5

6. Figure S5: Fitness surfaces with $\sigma_\vartheta$ = 2.5

7. Figure S6: Fitness surfaces with $\sigma_\vartheta$ = 3

8. Figure S7: Fitness surfaces with additive fitness effects, $\sigma_\vartheta$ = 0.5

9. Figure S8: Fitness surfaces with additive fitness effects, $\sigma_\vartheta$ = 2

10. Table S1: Exact maxima of fitness surfaces when $\sigma_\vartheta$ = 2

10. Table S2: Exact maxima of fitness surfaces when $\sigma_\vartheta$ = 0.5

11. Table S3: Exact maxima of fitness surfaces when $\sigma_\vartheta$ = 1

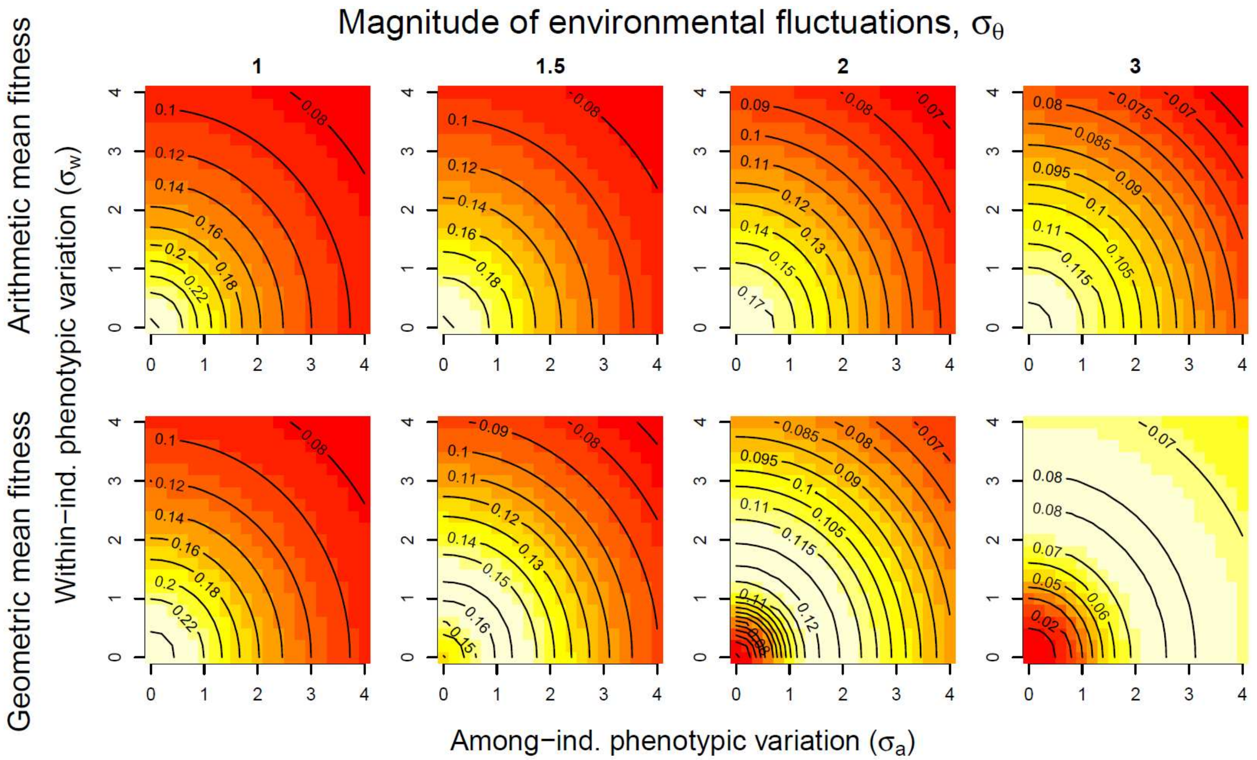


**Figure S1:** Fitness surfaces for genotypes with different amounts of among- ($\sigma_a$; x-axis) and within-individual ($\sigma_w$; y-axis) phenotypic variation. Contour lines show long-term arithmetic (top row) or geometric (bottom row) mean fitness in an environment where the phenotypic optimum $\Theta$ varies across generations according to a Gaussian distribution around 0 with standard deviation ± $\sigma_\Theta$ (values stated across the different columns). In this instance, there is no within-generation environmental variation ($\sigma_\vartheta$ = 0). Lighter background colors represent relatively higher fitness within each panel – note from the contour values that the different colors represent do not necessarily correspond to the same fitness values across panels.

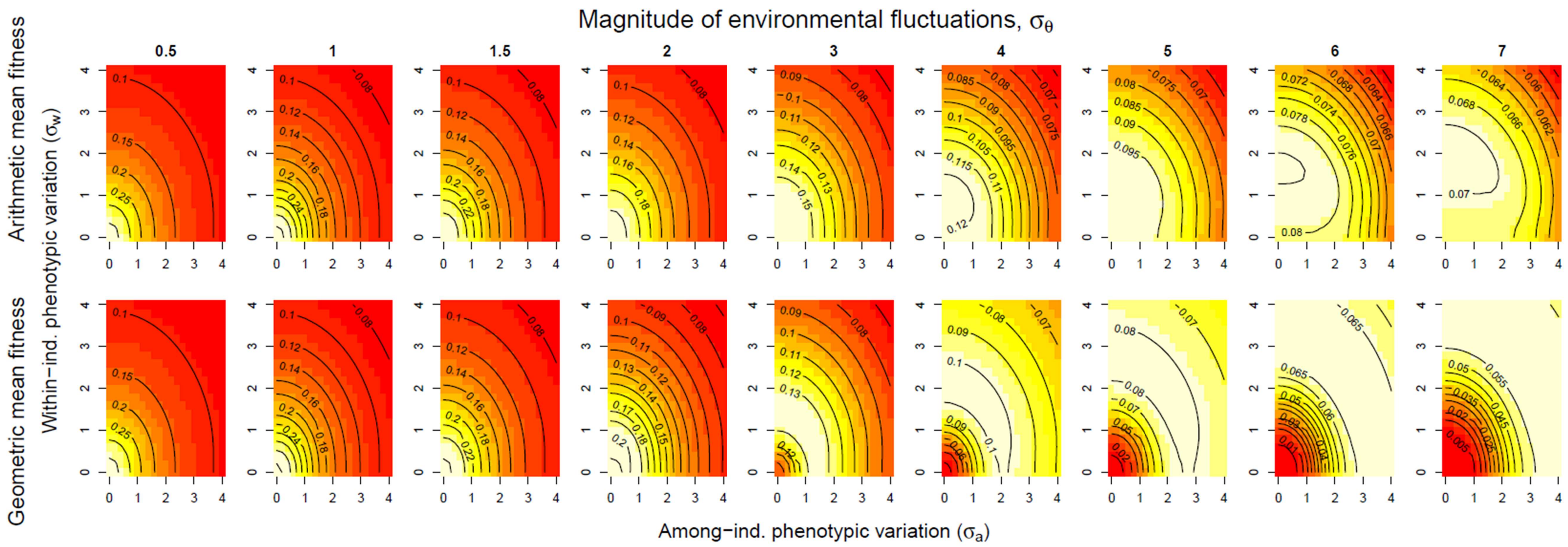


**Figure S2:** Fitness surface for genotypes with different amounts of among- ($\sigma_a$; x-axis) or within-individual ($\sigma_w$; y-axis) phenotypic variation. Contour lines show long-term arithmetic (top row) or geometric (bottom row) mean fitness in an environment where the mean optimal phenotype $\Theta$ varies across generations according to a uniform distribution around 0 with range ± $\sigma_\Theta$ (with values stated above the different columns). Within generations experienced environments $\vartheta$ vary temporally (with multiplicative fitness effects) according to a uniform distribution around $\Theta$ with range ± $\sigma_\vartheta$ = 0.5. Lighter background colors represent relatively higher fitness within each panel – note from the contour values that the different colors represent do not necessarily correspond to the same fitness values across panels.

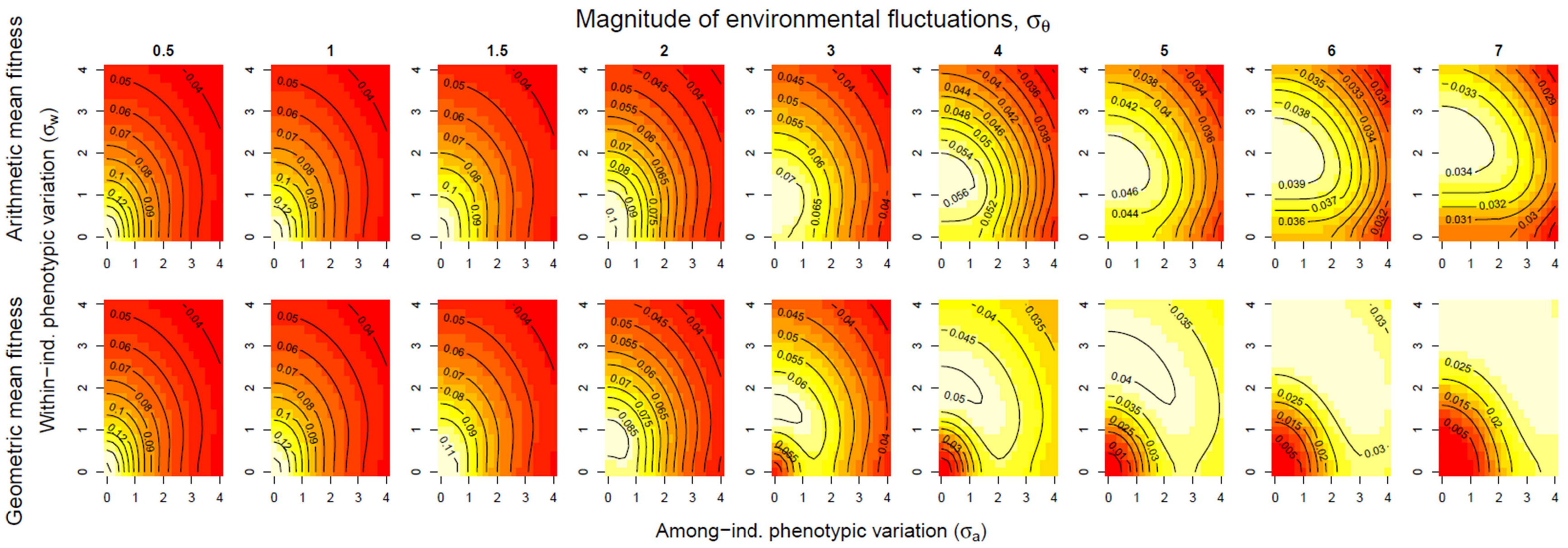


**Figure S3:** Fitness surface for genotypes with different amounts of among- ($\sigma_a$; x-axis) or within-individual ($\sigma_w$; y-axis) phenotypic variation. Contour lines show long-term arithmetic (top row) or geometric (bottom row) mean fitness in an environment where the mean optimal phenotype $\Theta$ varies across generations according to a uniform distribution around 0 with range $\pm\, \sigma_\Theta$ (with values stated above the different columns). Within generations experienced environments $\vartheta$ vary temporally (with multiplicative fitness effects) according to a uniform distribution around $\Theta$ with range $\pm\, \sigma_\vartheta = 1$. Lighter background colors represent relatively higher fitness within each panel – note from the contour values that the different colors represent do not necessarily correspond to the same fitness values across panels.

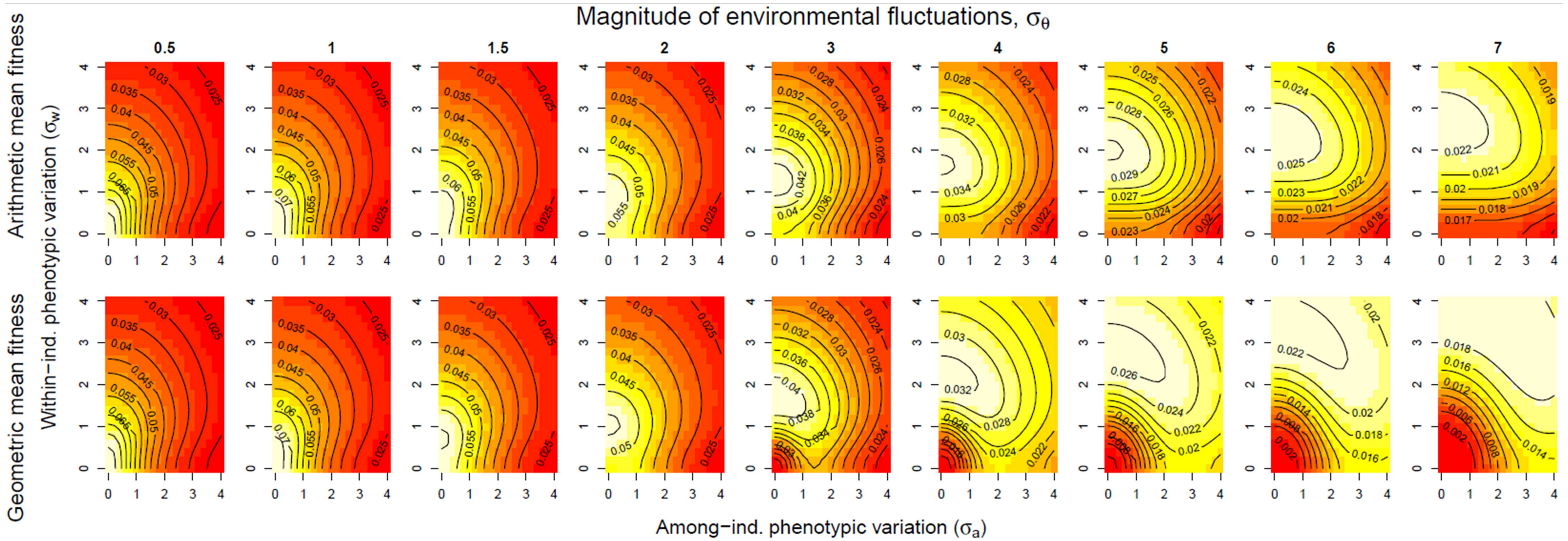


**Figure S4:** Fitness surface for genotypes with different amounts of among- ($\sigma_a$; x-axis) or within-individual ($\sigma_w$; y-axis) phenotypic variation. Contour lines show long-term arithmetic (top row) or geometric (bottom row) mean fitness in an environment where the mean optimal phenotype $\Theta$ varies across generations according to a uniform distribution around 0 with range ± $\sigma_\Theta$ (with values stated above the different columns). Within generations experienced environments $\vartheta$ vary temporally (with multiplicative fitness effects) according to a uniform distribution around $\Theta$ with range ± $\sigma_\vartheta$ = 1.5. Lighter background colors represent relatively higher fitness within each panel – note from the contour values that the different colors represent do not necessarily correspond to the same fitness values across panels.

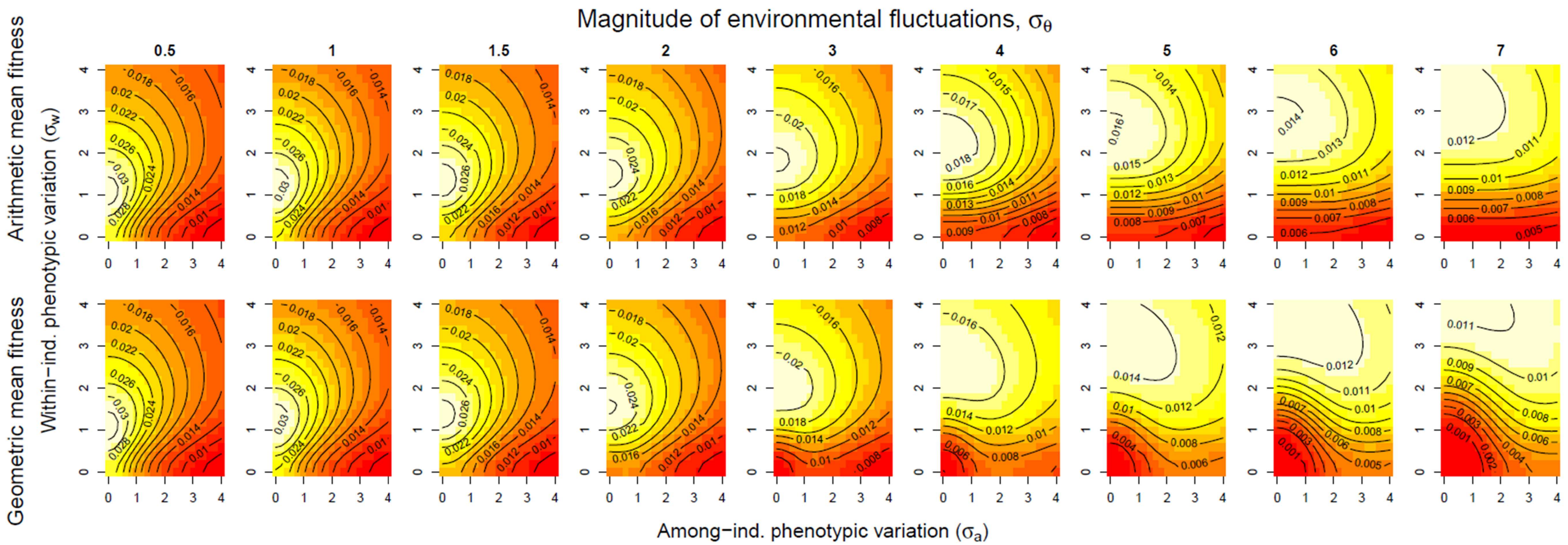


**Figure S5:** Fitness surface for genotypes with different amounts of among- ($\sigma_a$; x-axis) or within-individual ($\sigma_w$; y-axis) phenotypic variation. Contour lines show long-term arithmetic (top row) or geometric (bottom row) mean fitness in an environment where the mean optimal phenotype $\Theta$ varies across generations according to a uniform distribution around 0 with range ± $\sigma_\Theta$ (with values stated above the different columns). Within generations experienced environments $\vartheta$ vary temporally (with multiplicative fitness effects) according to a uniform distribution around $\Theta$ with range ± $\sigma_\vartheta$ = 2.5. Lighter background colors represent relatively higher fitness within each panel – note from the contour values that the different colors represent do not necessarily correspond to the same fitness values across panels.

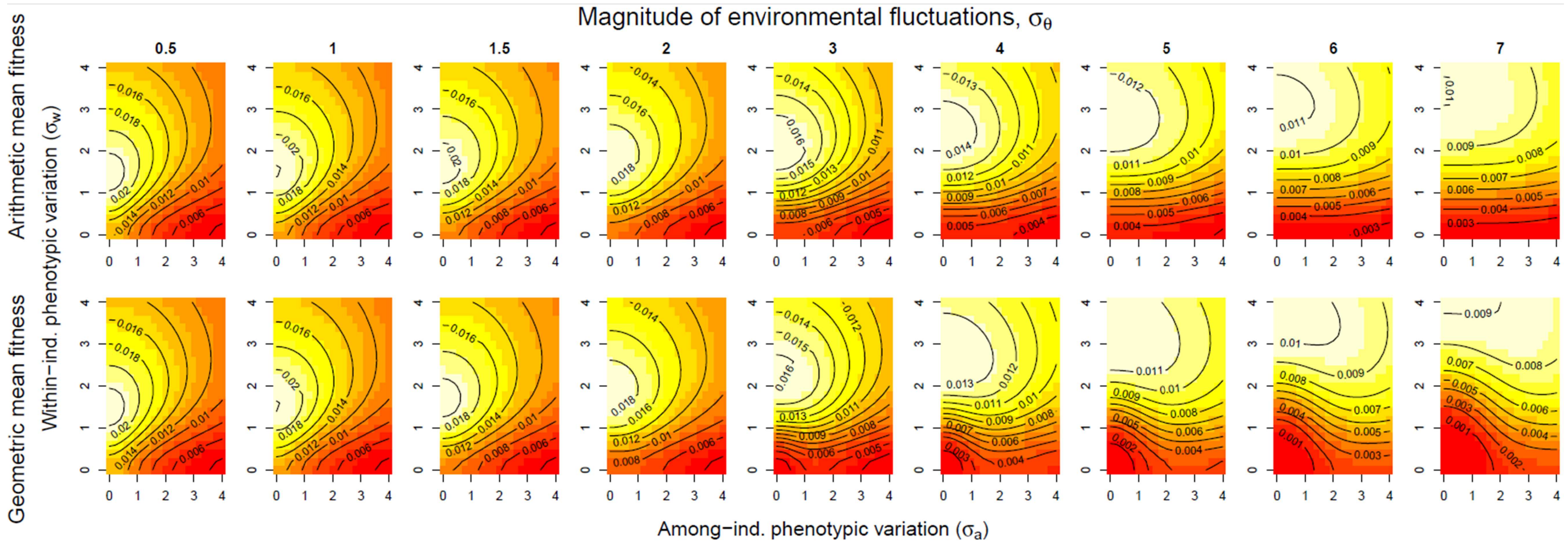


**Figure S6:** Fitness surface for genotypes with different amounts of among- ($\sigma_a$; x-axis) or within-individual ($\sigma_w$; y-axis) phenotypic variation. Contour lines show long-term arithmetic (top row) or geometric (bottom row) mean fitness in an environment where the mean optimal phenotype $\Theta$ varies across generations according to a uniform distribution around 0 with range $\pm\, \sigma_\Theta$ (with values stated above the different columns). Within generations experienced environments $\vartheta$ vary temporally (with multiplicative fitness effects) according to a uniform distribution around $\Theta$ with range $\pm\, \sigma_\vartheta = 3$. Lighter background colors represent relatively higher fitness within each panel – note from the contour values that the different colors represent do not necessarily correspond to the same fitness values across panels.

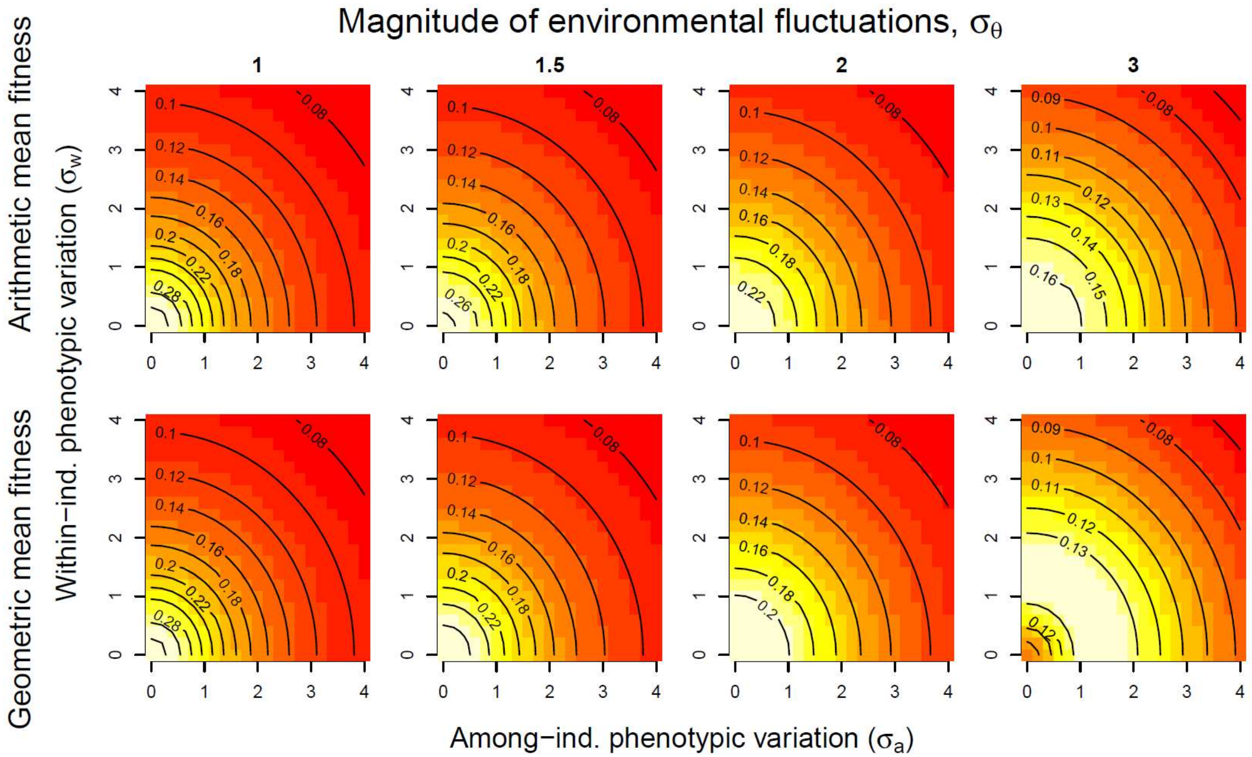


**Figure S7:** Fitness surface for genotypes with different amounts of among- ($\sigma_a$; x-axis) or within-individual ($\sigma_w$; y-axis) phenotypic variation. Contour lines show long-term arithmetic (top row) or geometric (bottom row) mean fitness in an environment where the mean optimal phenotype $\Theta$ varies across generations according to a uniform distribution around 0 with range ± $\sigma_\Theta$ (with values stated above the different columns). Within generations experienced environments ϑ vary temporally (with additive fitness effects) according to a uniform distribution around $\Theta$ with range ± $\sigma_\vartheta$ = 0.5. Lighter background colors represent relatively higher fitness within each panel – note from the contour values that the different colors represent do not necessarily correspond to the same fitness values across panels.

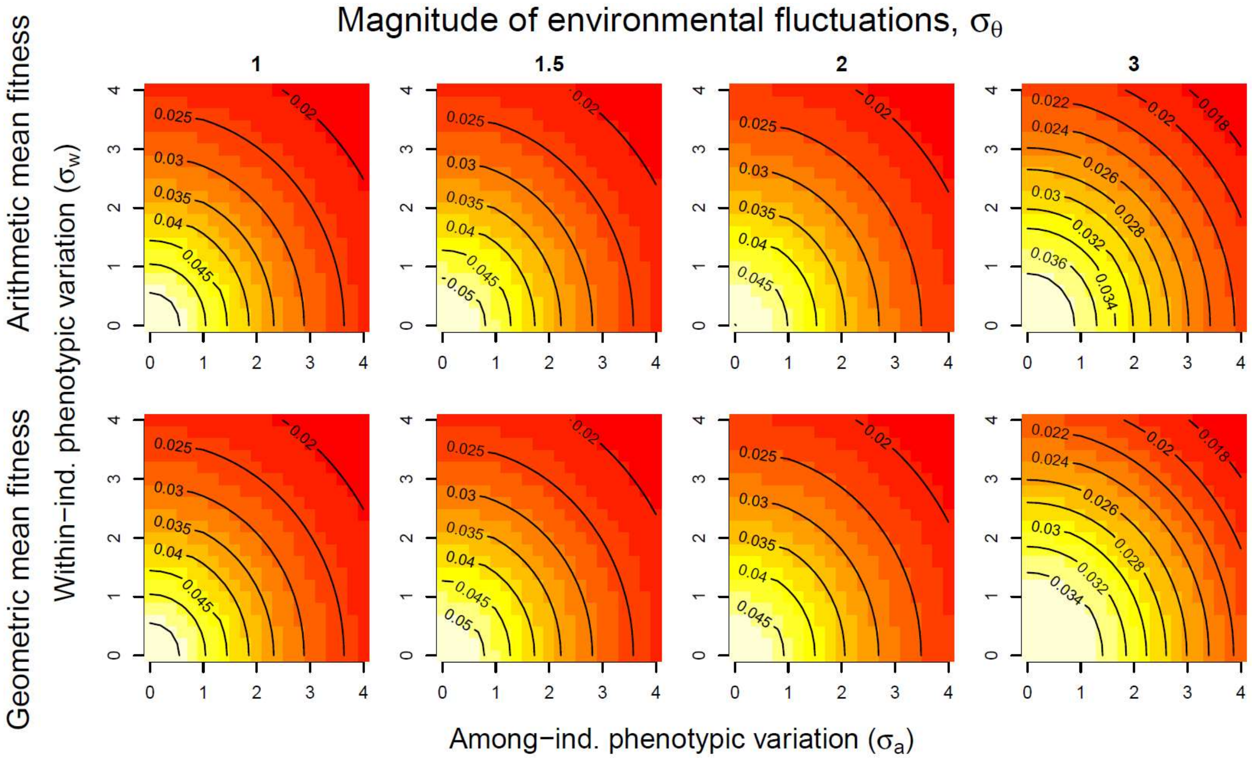


**Figure S8:** Fitness surface for genotypes with different amounts of among- ($\sigma_a$; x-axis) or within-individual ($\sigma_w$; y-axis) phenotypic variation. Contour lines show long-term arithmetic (top row) or geometric (bottom row) mean fitness in an environment where the mean optimal phenotype $\Theta$ varies across generations according to a uniform distribution around 0 with range ± $\sigma_\Theta$ (with values stated above the different columns). Within generations experienced environments $\vartheta$ vary temporally (with additive fitness effects) according to a uniform distribution around $\Theta$ with range ± $\sigma_\vartheta$ = 2. Lighter background colors represent relatively higher fitness within each panel – note from the contour values that the different colors represent do not necessarily correspond to the same fitness values across panels.

Table S1: Exact maxima of fitness surfaces when $\sigma_\vartheta$=2 (multiplicative within-generation fitness effects).

| $\sigma_\Theta$ | *Arithmetic mean fitness* | | *Geometric mean fitness* | | *Conservative bet-hedging* |
|---|---|---|---|---|---|
| | $\sigma_a$ | $\sigma_w$ | $\sigma_a$ | $\sigma_w$ | $\sigma_{w\text{ geom}} - \sigma_{w\text{ arit}}$ |
| 0.5 | 0 | 0.64 | 0 | 0.65 | 0.01 |
| 1 | 0 | 0.80 | 0 | 0.82 | 0.02 |
| 1.5 | 0 | 0.99 | 0 | 1.04 | 0.05 |
| 2 | 0 | 1.18 | 0 | 1.29 | 0.11 |
| 3 | 0 | 1.56 | 0 | 1.83 | 0.27 |
| 4 | 0 | 1.92 | 0 | 2.38 | 0.42 |
| 5 | 0 | 2.26 | 0 | 2.94 | 0.68 |
| 6 | 0 | 2.60 | 0 | 3.51 | 0.91 |
| 7 | 0 | 2.91 | 0 | 4.08 | 1.17 |

Table S2: Exact maxima of fitness surfaces when $\sigma_\vartheta$=0.5 (multiplicative within-generation fitness effects).

| $\sigma_\Theta$ | *Arithmetic mean fitness* | | *Geometric mean fitness* | | *CBH* |
|---|---|---|---|---|---|
| | $\sigma_a$ | $\sigma_w$ | $\sigma_a$ | $\sigma_w$ | $\sigma_{w\text{ geom}} - \sigma_{w\text{ arit}}$ |
| *0.5* | 0 | 0 | 0 | 0 | 0 |
| *1* | 0 | 0 | 0 | 0 | 0 |
| *1.5* | 0 | 0 | 0 | 0 | 0 |
| *2* | 0 | 0 | 0 | 0.65 | 0.65 |
| *3* | 0 | 0.54 | 0.02 | 1.44 | 0.90 |
| *4* | 0 | 1 | 0.02 | 2.10 | 1.10 |
| *5* | 0 | 1.36 | 0.23 | 2.71 | 1.37 |
| *6* | 0 | 1.67 | 0.23 | 3.32 | 1.65 |
| *7* | 0 | 2.03 | 0.23 | 3.92 | 1.89 |

Note in Table S2 that when within-generation environmental fluctuations $\sigma_\vartheta$ are very small relative to the between-generation fluctuations $\sigma_\Theta$, a small amount of DBH occurs at the optimum. This is also seen in that the geometric mean fitness landscape peaks at $\sigma_a > 0$ in the rightmost panels of Fig. S2, whereas arithmetic mean fitness landscapes always peak at $\sigma_a = 0$, indicating that DBH and CBH might coexist for only a very limited part of our parameter space under this scenario. However, this may also be one consequence of our use of Gaussian phenotypic distributions, where the heavy tails created by a large $\sigma_w$ are also able to deal with between-generation environmental fluctuations, whereas only when very low values of $\sigma_w$ are favored can DBH then also evolve on top.

Table S3: Exact maxima of fitness surfaces when $\sigma_\vartheta$=1 (multiplicative within-generation fitness effects).

| $\sigma_\Theta$ | *Arithmetic mean fitness* | | *Geometric mean fitness* | | *CBH* |
|---|---|---|---|---|---|
| | $\sigma_a$ | $\sigma_w$ | $\sigma_a$ | $\sigma_w$ | $\sigma_{w\,\mathrm{geom}} - \sigma_{w\,\mathrm{arit}}$ |
| *0.5* | 0 | 0 | 0 | 0 | 0 |
| *1* | 0 | 0 | 0 | 0 | 0 |
| *1.5* | 0 | 0 | 0 | 0.29 | 0.29 |
| *2* | 0 | 0.41 | 0 | 0.82 | 0.41 |
| *3* | 0 | 0.96 | 0 | 1.53 | 0.57 |
| *4* | 0 | 1.36 | 0 | 2.16 | 0.80 |
| *5* | 0 | 1.71 | 0 | 2.77 | 1.16 |
| *6* | 0 | 2.04 | 0 | 3.37 | 1.33 |
| *7* | 0 | 2.35 | 0 | 3.96 | 1.61 |
